# Targeting FEN1 to enhance efficacy of PARP inhibition in triple-negative breast cancer

**DOI:** 10.1101/2024.10.01.616136

**Authors:** Mallory I. Frederick, Elicia Fyle, Anna Clouvel, Djihane Abdesselam, Saima Hassan

**Affiliations:** Faculty of Medicine, Université de Montréal, Montréal, QC H3C 3T5, Canada; Centre de Recherche du Centre hospitalier de l’Université de Montréal (CRCHUM), l’Institut de Cancer de Montreal, Montreal, QC H2X 0A9, Canada; Division of Surgical Oncology, Department of Surgery, Centre hospitalier de l’Université de Montréal (CHUM), Montreal, QC H2X 0C1, Canada

**Keywords:** PARP inhibitor, FEN1 inhibition, triple-negative breast cancer, intrinsic and acquired resistance to PARP inhibitor, combination therapy, DNA damage, DNA replication fork speed

## Abstract

Patients with triple-negative breast cancer (TNBC) have limited targeted therapeutic options. PARP inhibitors (PARPi) have demonstrated an important role for *BRCA*-mutant patients with early TNBC. Combination approaches with PARPi can broaden the use of PARPi to a larger cohort of TNBC patients. We selected six genes from our previously identified 63-gene signature that was associated with PARPi response. siFEN1 increased cells in G2/M arrest, DNA damage and particularly apoptosis. Targeting FEN1 with a chemical inhibitor enhanced the efficacy of PARPi in 7/10 cell lines, and synergy was demonstrated mainly in PARPi-resistant TNBC cell lines. A *BRCA2*-mutant cell line with acquired resistance to olaparib (HCC1395-OlaR) was strongly synergistic, with a combination index value of 0.20. The combination of PARPi and FEN1 inhibition also showed synergy in a PARPi-resistant xenograft-derived organoid model. Two mechanisms which explain the underlying efficacy are rapid progression in DNA replication fork speed and enhancement of DNA damage. The combination induced the highest fork speed (47% difference in comparison to control, P<0.0001) when FEN1 inhibition and PARPi equally increased fork speed individually in a cell line with a pre-existing increase in replication stress. The combination also increased DNA damage at lower drug concentrations, driving response in most of the synergistic cell lines. Gene expression analysis suggested that the sensitizing role of FEN1 inhibition in PARPi-resistant cell lines may be due to downregulation of pathways including mismatch repair. Therefore, targeting FEN1 shows great therapeutic potential as a targeted combination approach, particularly in the context of PARPi-resistant TNBC.

## Introduction

Triple-negative breast cancer (TNBC) is the most aggressive subtype of breast cancer. TNBC makes up 15-20% of all breast cancers^1^ and has limited treatment options, primarily attributed to the lack of estrogen and progesterone hormone receptor (ER/PR) and human epidermal growth factor receptor 2 (HER2) overexpression. Among TNBC patients, 11-20% carry germline mutations in the *BRCA1*/*2* genes (gBRCA^MUT^)^1^, which has been associated with an increase in breast cancer risk^2^. In cells, BRCA1/2 function in the homologous recombination repair pathway to mend double strand breaks (DSBs) in DNA^3^. Loss of function of BRCA1/2, as in gBRCA^MUT^ cases, leads to improper repair of DSBs and contributes to the development of breast cancer.

Patients who are gBRCA^MUT^ benefit from Poly (ADP-ribose) polymerase (PARP) inhibitors (PARPi), which act by synthetic lethality and PARP-DNA trapping^4^. PARPi are the only targeted therapy available for early TNBC patients, including talazoparib and olaparib. Talazoparib is the most potent in the context of PARP-DNA trapping^4^. However, studies have shown that PARPi may also be effective in tumors that actually lack the *BRCA1/2*-mutation, yet demonstrate ‘BRCAness’, with a functional defect in DNA repair commonly attributed to impaired homologous recombination^5,6^. Several biomarkers of BRCAness have been tested in patient samples including those derived from whole genome sequencing (HRDetect), genomic scar assays, gene expression signatures, mutational signatures, and detection of RAD51 foci by immunofluorescence^1^.

We previously derived a 63-gene signature predictive of PARPi response in TNBC^7^. Using the DNA damage response of three PARPi in a panel of TNBC cell lines, we showed that PARPi response was associated with an enrichment of pathways including DNA double strand break repair, DNA damage bypass, base excision repair, and cell cycle checkpoints. Our gene signature demonstrated an overall accuracy of 86% in a small cohort of patient-derived xenografts treated with olaparib, and predicted sensitivity in 45% of TNBC patients, which was comparable to other biomarkers of BRCAness^1^. However, in addition to being a predictive biomarker, our gene signature can also be used to identify therapeutic targets^7^. These targets can enhance the activity of anti-PARP therapy by targeting pathways involved in DNA repair or cell cycle checkpoints.

Clinically promising therapeutic targets of genes from our gene signature have emerged. For example, *CHK1*, involved in checkpoint mediated cell cycle arrest, and *USP1*, ubiquitin specific peptidase 1, involved in DNA damage bypass, have demonstrated efficacy when targeted in combination with PARPi preclinically^8,9^, and are being tested in Phase 1 clinical trials in solid tumors^10,11^. PARPi have also been combined with inhibitors of *ATM*, a DNA damage kinase, or WEE1, a regulator of cyclin-dependent kinase 1/2^12-14^. Hence, there is precedence to combine PARPi with drugs that target DNA repair or cell cycle checkpoints^15^.

Here, we screened targets from the 63-gene signature and selected Flap Endonuclease 1 (FEN1) as a target for PARPi combination therapy. FEN1 is a replication-regulating protein, involved in base excision repair, and is recruited to single-strand DNA damage sites and Okazaki sites by active PARP-1^16,17^. Interestingly, FEN1 has been identified in other PARPi sensitivity screens^18,19^. Here, we used the FEN1 inhibitor LNT1^20^ in chemosensitivity assays to test its efficacy in combination with the PARPi talazoparib in TNBC cell lines and xenograft-derived organoid (XDO) models. We show that talazoparib is synergistic with LNT1 in TNBC cell lines with intrinsic and acquired resistance to PARPi. These cell lines demonstrated a marked increase in DNA fork replication speed and DNA damage. Together, these data indicate FEN1 as a promising target for use in PARPi combination therapy.

## Materials and methods

### Cell culture

A complete list of cell lines, sources, and corresponding media is available in Supplementary Table S1. Cell lines were validated by DNA fingerprinting using short-tandem repeat (STR) analysis, last performed in April 2024. Cell lines and organoids were tested regularly to confirm the absence of mycoplasma using Mycoalert mycoplasma detection kit (LT07, Lonza).

### siRNA transfections

We used ON-TARGETplus siRNA SMARTpools (Dharmacon) for human *BRCA1* (as a positive control), *BARD1, BUB1, RRM2, USP1, FEN1*, and *EXO1*, and non-target siRNA control (Dharmacon). Reverse transfection with Lipofectamine RNAimax (Invitrogen) was performed, with further details provided in Supplementary Methods.

### Acquired resistance cell line derivation

PARPi-resistant HCC1395 cells were derived from HCC1395 TNBC cells grown in RPMI with increasing concentrations of talazoparib or olaparib added to the media over the course of six months, as previously described^21^. Cells were passaged in T-25 flasks and allowed to adhere. The next day, media containing 0.0006 µM talazoparib or 0.14 µM olaparib was added. Cells were grown in the presence of either drug until confluence was reduced to approximately 10%, then allowed to recover to near confluence in complete media lacking PARPi. Media was exchanged every 3-5 days, alternating between media containing or lacking PARPi depending on cell growth. Treatment with PARPi was repeated at the same concentration until resistance was observed, at which point PARPi concentration was doubled. This process was repeated until sufficient resistance was observed. Finally, stable resistance was confirmed by IC_50_ calculation after 5 passages in the absence of PARPi and a freeze-thaw cycle.

### Western blots

Cells were lysed in a cell lysis buffer (Cat. #9803, Cell Signaling Technology) containing phosphatase inhibitors (Cat. #4906845001, Roche) and PMSF (Cat. #8553S, Cell Signaling Technology), followed by centrifugation at 14,000 x g for 10 minutes at 4°C. Protein concentrations were quantified using the Pierce™ BSA Protein Assay (Cat. #23227, Thermofisher). Equal amounts of protein were run on 8-12% acrylamide gels at 90 V for 30 minutes and 120 V for 1-2 hours, then transferred to nitrocellulose membranes for 1 hour at 100 V. Membranes were blocked with 5% BSA or milk in TBST buffer for 1 hour at room temperature, then incubated with primary antibodies overnight at 4°C. Membranes were washed 3 × 10 minutes in TBST buffer and incubated with the appropriate HRP-conjugated secondary antibodies for 1-2 hours at room temperature. Finally, membranes were washed 3 × 5 minutes in TBST, stored in TBS, and exposed to Clarity ECL (Biorad) or PICO ECL supersignal (ThermoFisher) to be imaged with the ChemiDoc Imaging System. All antibodies are listed in Supplementary Table S2.

### Xenograft-derived organoids

Organoids were developed from mice tumors obtained from orthotopic xenograft experiments that we previously performed^22^. All animal experiments were approved by the Institutional Animal Protection Committee (CIPA) of the Centre de Recherche de Centre hospitalier de l’Université de Montréal (CRCHUM) under protocol C17017SHs. Two million HCC1806 cells were surgically implanted in the mammary fat pad of NOD-SCID gamma (NSG) female mice (Cat. #005557, Jackson Laboratory). Once untreated tumors reached a mean volume of 12 mm^2^, mice were sacrificed and tumors harvested.

Fresh tumor samples were processed and organoid models established as described^23^. XDO maintenance media was adapted from Type 1 medium^23^ lacking primocin, with the following adjustments: 100 ng/mL R-spondin 1 (Cat. #120-38-100UG, PeproTech), 100 ng/mL Noggin (Cat. #120-10C-100UG, PeproTech), 5 mM nicotinamide (Cat. #A15970.22, Thermo Scientific), and 500 nM SB202190 (Cat #202190, StemCell Technologies). For 10-day chemosensitivity assays, 96-well plates (Cat. #3610, Corning) were pre-coated with 10 µL of Cultrex Reduced Growth Factor Basement Membrane Extract (Cat. #3433-005-01, R&D Biosystems). XDOs were trypsinized to single cells, resuspended at 20 000 cells/mL, and 100 µL of cell suspension was plated per well. Media lacking N-acetylcysteine and Y-27632 was used for XDO chemosensitivity assays, as previously performed^24^. XDOs were treated with talazoparib, LNT1, or the combination as described above for a total of 10 days. Cell viability was measured by CellTiter-Glo 3D Cell Viability Assay (Cat. #G9682, Promega) according to manufacturer protocols.

### RNA-seq analysis

RNA sequencing was performed on 3 independent samples collected from each cell line. RNA was extracted using RNeasy Plus mini kit (Cat. #74134). RNA was quantified using Qubit (Thermo Scientific) and quality was assessed with the 2100 Bioanalyzer (Agilent Technologies). Libraries were prepared and samples were sequenced with Illumina NextSeq 500, using the Illumina kit NextSeq 75 cycles High Output v2, with ∼ 25M reads per sample, at the Institute for Research in Immunology and Cancer (IRIC, Montreal).

The first part of the analysis including normalization and DeSEq2 analysis was performed at the Bioinformatics Facility at IRIC. Sequences were trimmed for sequencing adapters and low quality 3’ bases using Trimmomatic version 0.35^25^ and aligned to the reference human genome version GRCh38 (gene annotation from Gencode version 37, based on Ensembl 103) using STAR version 2.7.1a^26^. DESeq2 version 1.30.1^27^ was then used to normalize gene readcounts and produce the sample clustering. Using the log2-fold change values from the DESeq2 analysis, all expressed genes were ranked and a pre-ranked Gene Set Enrichment Analysis was performed using GenePattern^28^. Gene sets from MsigDB were used, including Hallmarks and KEGG legacy for human tissue. FDR cutoff of 25% and p-value < 0.05 were used to identify statistically significant gene sets. Dot plots and heatmaps of hierarchical clustering were created in RStudio v4.2.2^29^.

### Statistical analysis

Graphs with statistical analysis were made in GraphPad Prism 10, with the exception of an online tool, gene expression-based outcome for breast cancer, for clinical correlations^30^. Experiments were performed with a minimum of n=2 independent replicates, with 3 replicate wells each for experiments performed in 96-well plates. Significance was calculated by one- or two-way ANOVA with Tukey HSD as appropriate, apart from DNA fiber assays, which were analysed by Kruskal-Wallis ANOVA with multiple comparisons. P<0.05 is considered statistically significant.

Further details regarding siRNA transfections, chemosensitivity assays for IC_50_ and combination index values, DNA fiber assays, and flow cytometry are provided in Supplementary Methods^7,22^.

## Results

### Knockdown of selected genes from 63-gene signature increases cellular response to talazoparib

A 63-gene signature of PARPi response was previously derived from a panel of TNBC cell lines treated with veliparib, olaparib, or talazoparib^7^. We selected talazoparib for this study since this is a potent PARPi^4^, with similar pre-clinical and clinical efficacy as olaparib. From the signature, we identified seven genes of interest: *BRCA1*, BRCA1-associated RING domain protein 1 (*BARD1*), Mitotic checkpoint serine/threonine-protein kinase (*BUB1*), Ribonucleotide reductase regulatory subunit M2 (*RRM2*), *USP1, FEN1*, and Exonuclease 1 (*EXO1*). Targeted gene silencing using siRNA knockdown was performed in a *BRCA1/2*-wild type (BRCA^WT^) cell line, MDAMB231, which was known to overexpress each of these proteins (Fig. 1A). To determine whether targeting these proteins would increase the efficacy of talazoparib treatment, we used flow cytometry to quantify cell cycle changes, the proportion of γ-H2AX-positive (γ-H2AX+) cells, a marker of DSBs, and cleaved-Caspase 3-positive (cl-Caspase 3+) cells, a marker of apoptosis (Fig.1B-E).

**Figure 1.**
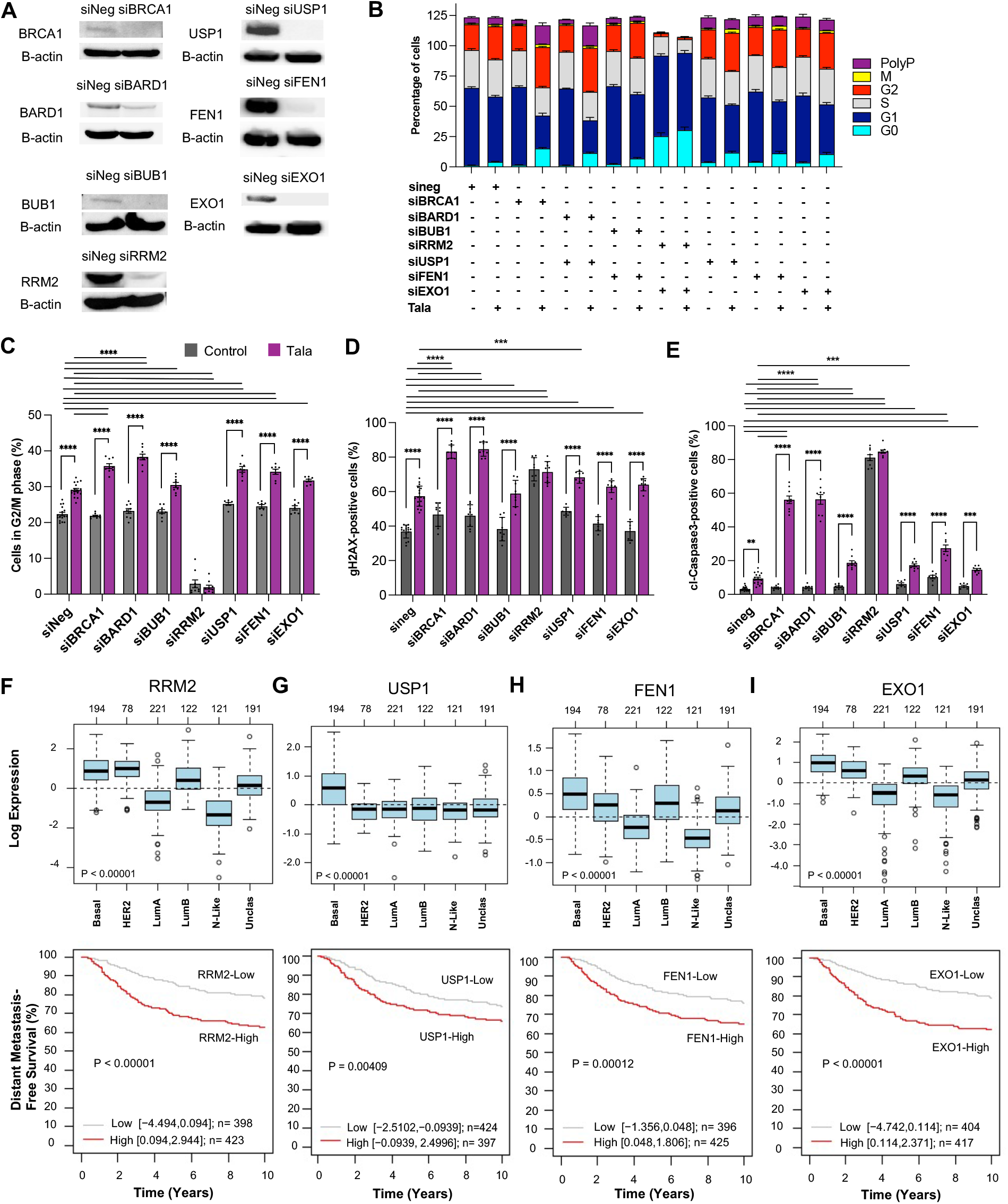
siRNA knockdown of selected genes from 63-gene signature induces G2/M arrest, DNA ge and apoptosis. siRNA knockdown of *BRCA1, BARD1, BUB1, RRM2, USP1, FEN1*, and *EXO1* in B231 TNBC cells with (A) Immunoblot validation; and 72-hour treatment in the absence or presence µM talazoparib with flow cytometry analysis for (B) Cell cycle analysis, (C) G2/M phase distribution, H2AX expression, and (E) Cleaved-caspase 3 expression. 2-way ANOVA and multiple comparisons erformed in GraphPad Prism. Data represent mean with SEM bars from three replicate experiments. ANOVA with Tukey’s multiple comparison test was performed. P<0.0001****, P<0.001***, P<0.01**. ohort of 881 untreated breast cancer patients (F-I), box plots evaluating gene expression across 0 subtypes along top panel and Kaplan-Meier survival curves for distant metastasis-free survival with r censoring along bottom panel^30^. Using Affymetrix U133A microarrays, gene expression is shown RRM2, (G) USP1, (H) FEN1, and (I) EXO1. For the top panel, PAM50 subtypes include basal, HER2, al A (LumA), Luminal B (LumB), Normal-Like (N-Like), and Unclassified (Unclass). P-values represent ANOVA test. For the bottom panel, Logrank P-values are shown for the survival analysis.

The impact of siBARD1 plus talazoparib was very similar to what was observed with siBRCA1, with an increase in cells in G2/M arrest (Δ9.2 or 15.9%), γ-H2AX+cells (Δ27.3 or 47.8%), and cl-Caspase 3+ cells (Δ47.0 or 53.1%) in comparison to talazoparib alone or control (P<0.0001 for each comparison). Combining talazoparib with either siBUB1 or siEXO1 resulted in similar increases in the percentage of cells in G2/M arrest (Δ8.1% or 9.4%), γ-H2AX+ cells (Δ22.0% or 27.2%), and cl-Caspase 3+ cells (Δ15.5% or 11.4%) in comparison to control (P<0.0001 for each comparison). However, neither the siBUB1 or siEXO1 combination with talazoparib had an impact on G2/M arrest or DNA damage in comparison to talazoparib alone. Each gene knockdown demonstrated an enhanced effect in combination with talazoparib in comparison to its respective gene knockdown alone, except for siRRM2. siRRM2 alone demonstrated a unique profile, leading to an increase in G0 and G1 cells, and was sufficient to increase DNA damage (Δ36.1% γ-H2AX+, P<0.0001) and apoptosis (Δ77.9% cl-Caspase 3+, P<0.0001). No additional benefit of talazoparib was also observed when combined with a small molecule inhibitor of RRM2, TAS1553^31^ (Supplementary Fig. S1).

Knockdown of two gene targets induced important cell cycle changes, DNA damage, and cell death: *USP1* and *FEN1*. The combination of talazoparib with siUSP1 or siFEN1 showed statistically significant increases in G2/M arrest and apoptosis in comparison to control and talazoparib alone. In particular, siUSP1 plus talazoparib increased the percentage of cells in G2/M arrest by 12.5% and 5.8% (P<0.0001; P<0.0001), and cl-Caspase 3+ cells by 14.0% and 7.9% (P<0.0001; P=0.0005), in comparison to control and talazoparib, respectively. siFEN1 plus talazoparib augmented the percentage of cells in G2M arrest by 11.8% and 5.1% (P<0.0001; P<0.0001) and demonstrated an important increase in cl-Caspase 3+ cells by 24.2% and 18.0% (P<0.0001; P<0.0001), in comparison to control and talazoparib, respectively.

To better understand the clinical significance of these targets, we evaluated the expression of these genes in different breast cancer subtypes a cohort of 881 untreated breast cancer patients^30^. We identified elevated expression of *USP1, RRM2, FEN1*, and *EXO1* in more aggressive breast cancer subtypes (basal and HER2), and that their overexpression were associated with a poorer 10-year distant metastasis-free survival (Fig. 1F-I). Of note, no statistical significance in distant metastasis-free survival was identified for *BARD1* or *BUB1* (data not shown).

To summarize, targeting genes from our 63-gene signature can induce greater cell sensitivity in combination with talazoparib. Focusing on those targets that have a commercially available inhibitor (BUB1, USP1, and FEN1), only USP1 and FEN1 were strong prognostic biomarkers. Since the greatest induction of cell death was demonstrated with siFEN1 plus talazoparib in comparison to control or talazoparib, we selected FEN1 for further investigation, using LNT1 as the chemical inhibitor^20^.

### LNT1 is synergistic with talazoparib in PARPi-resistant cell lines

To compare the impact of LNT1 with siFEN1, we evaluated cell cycle changes in MDAMB231 (Supplementary Fig. S1). The combination of LNT1 plus talazoparib increased the percentage of cells in G2/M arrest by 11%, which was similar to the proportion of cells in G2/M arrest induced by siFEN1 and talazoparib (Fig. 1C).

First, we determined IC_50_ values for the single agents LNT1 and talazoparib in ten TNBC cell lines, including two with acquired resistance to PARPi. Molecular features of these cell lines have been previously summarized^22^. We developed two cell lines with acquired resistance to PARPi by exposing talazoparib-sensitive HCC1395 TNBC cells (*BRCA2*-mutant) to increasing concentrations of olaparib (olaparib-resistant, OlaR) or talazoparib (talazoparib-resistant, TalaR) (Fig. 2A). Both resistant cell lines reached resistance to talazoparib in a similar manner to our most intrinsically resistant cell line, BT549 (BRCA^WT^) (Fig. 2B). The IC_50_ values for talazoparib among our cell lines ranged from 3.0 × 10^−4^ µM to 0.47 µM, while LNT1 sensitivity was in a much smaller range, from 1.4 µM to 15.7 µM (Fig. 2B, C). Similar responses to PARPi and LNT1 were demonstrated in some cell lines, with MX1 (*BRCA1*-deleted and *BRCA2*-mutant), MDAMB436 (*BRCA1*-mutant), and HCC1806 (BRCA^WT^) showing sensitivity, while HCC1395-OlaR, HCC1395-TalaR, and Hs578T showing resistance to each of the compounds. However, some cell lines had different responses, with SUM149PT (*BRCA1*-mutant) being the most sensitive to LNT1 with intermediate sensitivity to talazoparib, and BT549 having intermediate sensitivity to LNT. Therefore, *BRCA1/2*-mutation status is not required for LNT1 sensitivity.

**Figure 2.**
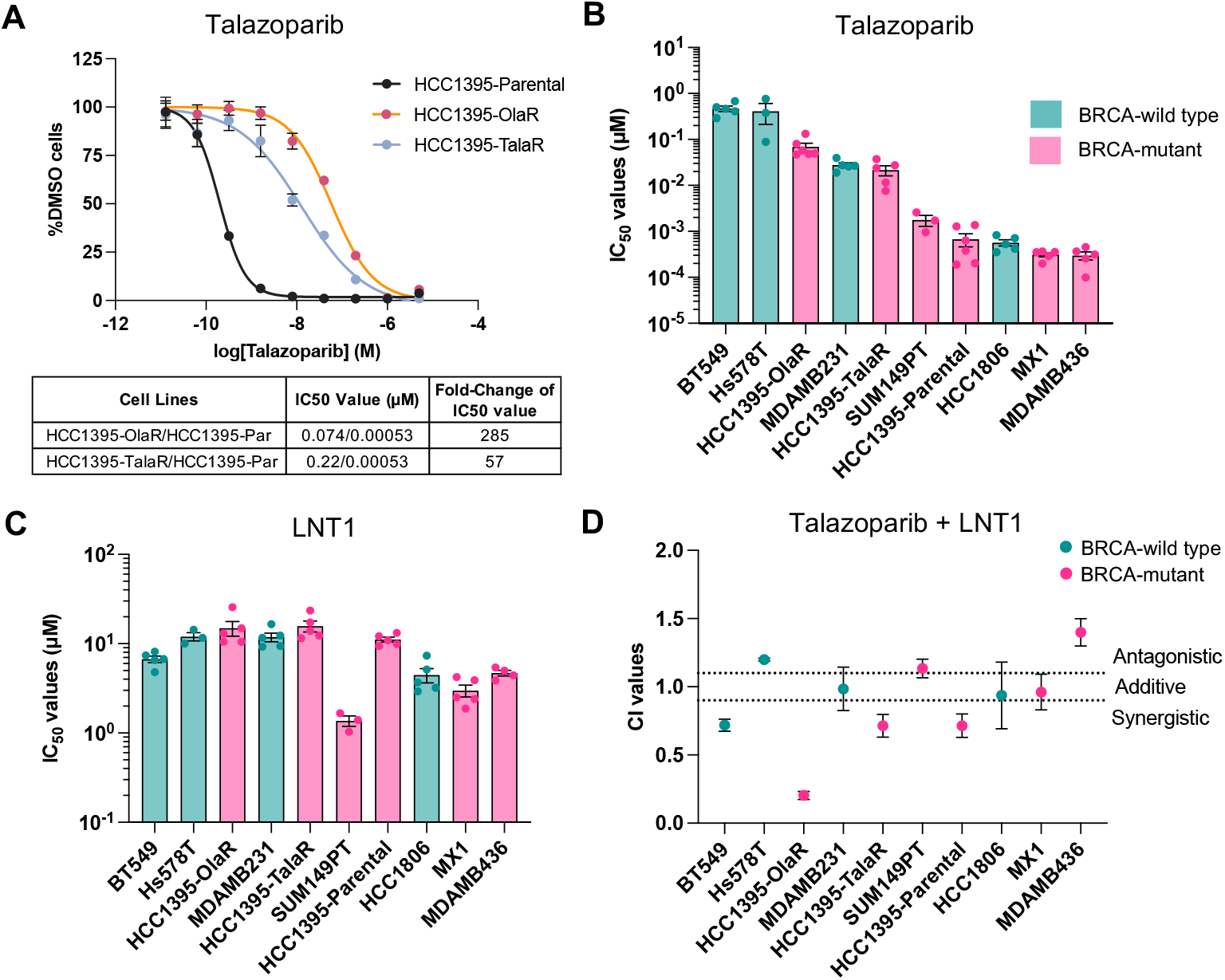
LNT1 enhances sensitivity to PARPi in most TNBC cell lines. HCC1395 cell lines were d with acquired resistance to PARP inhibitors. (A) IC_50_ dose-response curves for HCC1395-Parental 1395-Par) (black), HCC1395-Olaparib Resistant (OlaR) (orange), and HCC1395-Talazoparib tant (TalaR) (blue) cell lines. IC_50_ values for the resistant cell lines and ratio relative to HCC1395-tal are indicated in table below. IC_50_ values for (B) talazoparib, and (C) LNT1 in TNBC cell lines. Data sent mean + /− SEM, with individual replicates indicated. (D) Combination index values for talazoparib NT1 reported at an FA of 0.5. Teal green bars/dots represent BRCA^WT^ cell lines, and pink bars/dots sent BRCA^MUT^ cell lines.

Next, we determined the efficacy of the combination of LNT1 and talazoparib by calculating Combination Index (CI) values. Here, we considered CI < 0.9 to be synergistic, 0.9 < CI < 1.1 as additive, and CI > 1.1 as antagonistic^32^. Overall, four TNBC cell lines were synergistic, three were additive, and three were effectively antagonistic (Fig. 2D). SUM149PT was borderline antagonistic, with a CI value of 1.1, while Hs578T and MDAMB436 had CI values of 1.2 and 1.4, respectively. Surprisingly, the cell lines that were highly resistant to talazoparib showed the highest synergy with LNT1, including BT549 (0.7), HCC1395-OlaR (0.2), and HCC1395-TalaR (0.7). Among these three cell lines, the dose reduction index of talazoparib ranged from 3.4 to 14.7 and from 1.8 to 7.5 for LNT1. This indicates that, to achieve synergy in these PARPi resistant cell lines, the levels of LNT1 and talazoparib required can be drastically reduced while still inducing cell death.

### Talazoparib and LNT1 induce DNA damage and apoptosis in PARPi-resistant cell lines

To investigate the mechanism of synergy between talazoparib and LNT1 in talazoparib-resistant cell lines, we measured induction of DNA damage and apoptosis by immunofluorescence. Cells were treated with increasing concentrations of LNT1 (L1-L6), talazoparib (T1-T6), or the combination (TL1-TL6) for ten days (Supplementary Table S3), then fixed and stained for 53BP1 and cleaved-PARP. We calculated a 53BP1 product score for DNA damage using the product of the mean number of nuclear 53BP1 foci per cell and the percentage of cells positive for 53BP1 (Fig. 3, Supplementary Fig. S3)^22^. We found that, in talazoparib-resistant cells, the combination of LNT1 with talazoparib induced DNA damage at lower concentrations than either agent alone (Fig. 3A). In HCC1395-OlaR cells, the most synergistic cell line, we observed DNA damage at TL3, which was similar to the levels observed with T5 and L5 treatments alone (Fig. 3A). A similar trend was observed in HCC1395-TalaR cells, where ubiquitous DNA damage was observed at TL4, with much lower levels at T6 and similar levels at L5. In contrast, we performed the same experiment in cell lines that exhibited near-additive (MDAMB231, HCC1806) CI values for the talazoparib and LNT1 combination and found a much less pronounced effect from the combination, with DNA damage induced at similar concentrations in the combination as single agent treatments, albeit at different levels.

**Figure 3.**
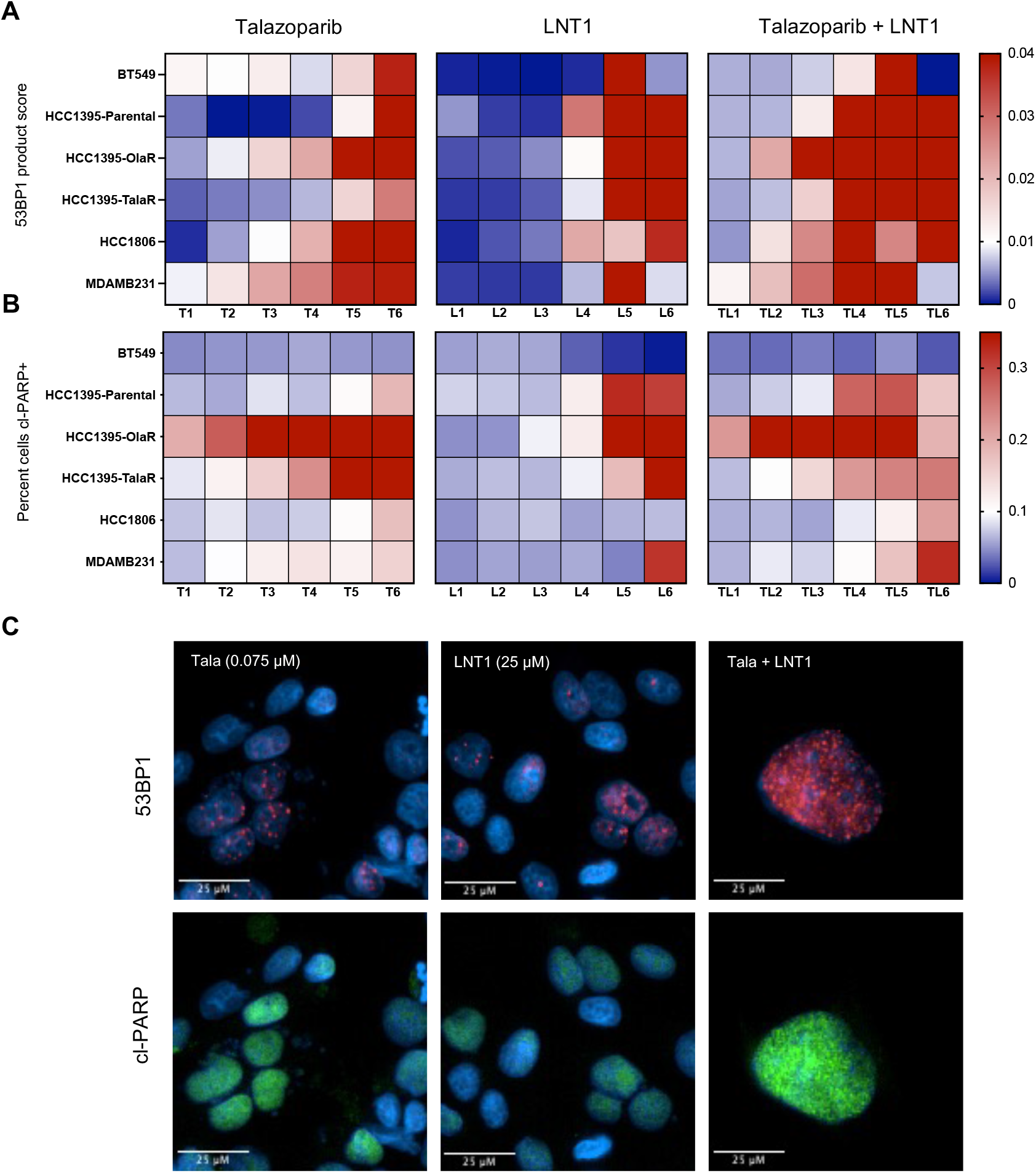
Combined talazoparib and LNT1 induces DNA damage at reduced concentrations in rgistic cell lines. Heatmaps of normalized (A) 53BP1 product score (product of mean 53BP1 foci per and percentage of cells positive for 53BP1) and (B) percentage of cleaved-PARP positive (cl-PARP+) The x-axis indicates increasing concentrations (1-6) of talazoparib (T or Tala), LNT1 (L), or the ination (TL), with concentrations centered around the IC_50_ for each cell line. (C) Representative 20x tive images of HCC1395-OlaR cells treated with talazoparib (left), LNT1 (middle), or the combination Cells were stained for 53BP1 foci (red) and cl-PARP+ cells (green), with nuclear staining in blue. bars indicate 25 µm.

Apoptosis was measured by the percent of cells positive for cleaved-PARP (cl-PARP+) Fig. 3B)^22^. As we had seen a more pronounced effect of FEN1 knockdown on apoptosis than induction of DNA damage (Fig. 1), we here expected a similar trend. Indeed, in the synergistic HCC1395-parental cell line, we saw increased induction of apoptosis at TL4, while similar levels were induced at T6 and L5 in the individual treatments. As with DNA damage, a less pronounced effect on apoptosis was observed in the two additive cell line, HCC1806 and MDAMB231, where the percent of cl-PARP+ cells was similar in combination as to talazoparib or LNT1 alone. Interestingly, the highly synergistic OlaR cells exhibited high levels of cl-PARP+ cells even at low concentrations of talazoparib alone (Fig. 3B). This may indicate that OlaR cells are adapted to endure a high level of DNA damage, as induced by talazoparib treatment.

### Combined PARPi and LNT1 increases DNA fork speed in replicating cells

Increases in replication fork speed in response to PARPi and FEN1 depletion have previously been reported^16,22^. Therefore, we investigated the impact of the combination of 72h treatments with talazoparib and/or LNT1 on fork replication speed in four TNBC cell lines (Fig. 4, Supplementary Fig. S4). We observed differences in endogenous replication speed depending on the cell line, but consistent increases in fork speed in talazoparib treated cells. In contrast, LNT1 increased the fork speed by 26% (P<0.0001) in only one cell line BT549, a synergistic cell line (Fig. 4A). Interestingly, the highest increase in fork speed induced by the combination was also demonstrated in BT549 in comparison to control (47.4%, P<0.0001), and ∼17% increase in comparison to either talazoparib alone (P=0.008) or LNT1 alone (P=0.0002). In the most synergistic cell line, HCC1395-OlaR, the combination increased fork speed by 30.9% in comparison to control (P<0.0001) and 26.1% in comparison to talazoparib alone (P<0.0001) (Fig. 4B). In MDAMB231, which demonstrated a nearly additive effect of the combination, fork speed increased by 23.6% in comparison to control (P<0.0001) (Fig. 4C). Finally, another cell line which also showed a nearly additive effect with the combination, MX1, revealed an increase in fork speed by 25.0% in comparison to control (P<0.0001) and a 10.0% increase in comparison to talazoparib (P=0.002) (Fig. 4D). Overall, since high fork speeds were more pronounced in the synergistic cell lines, acceleration of fork speed is an important mechanism explaining the efficacy of the combination of talazoparib and LNT1.

**Figure 4.**
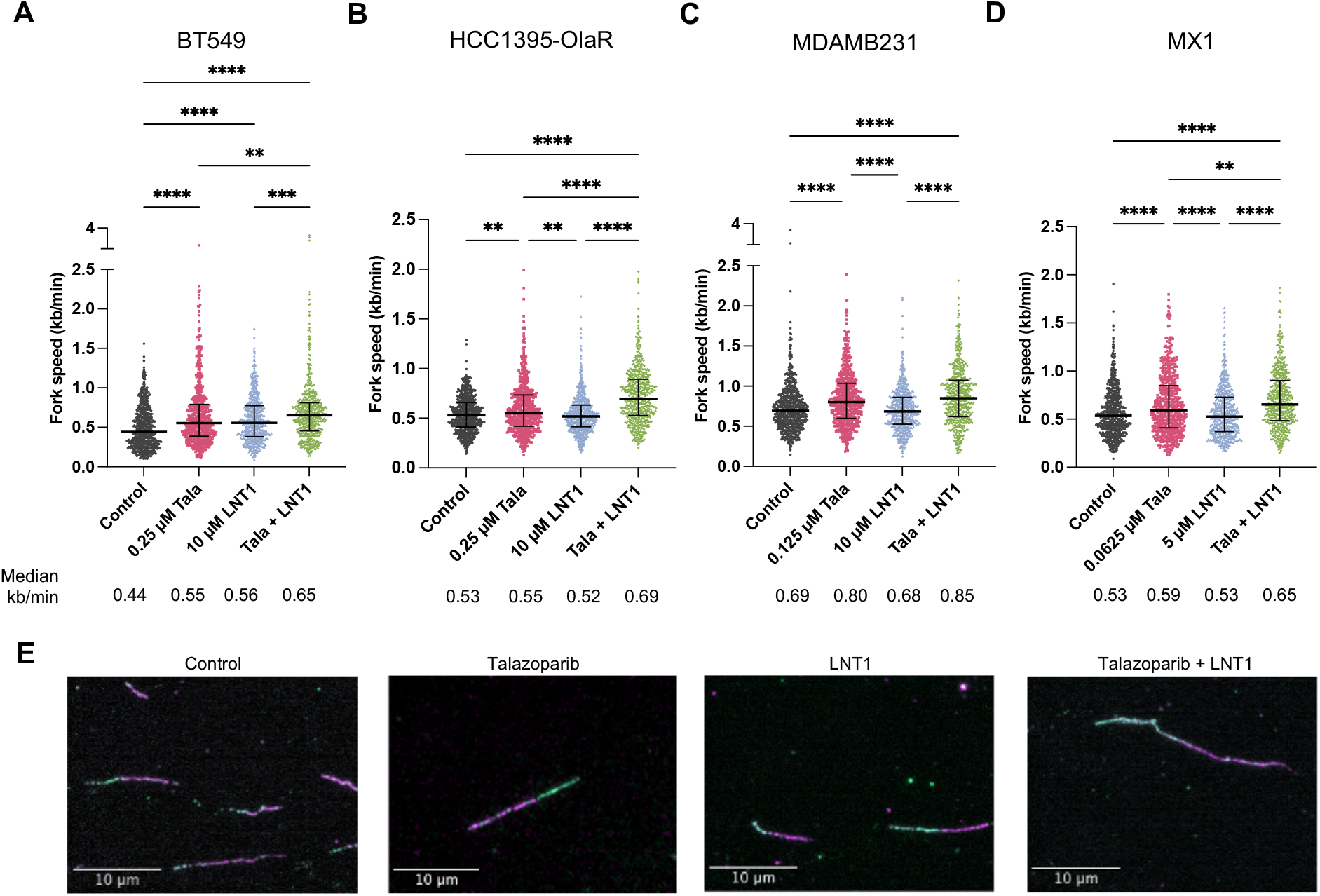
Talazoparib and LNT1 increase DNA replication fork speed in TNBC cells. (A-D) DNA fork for (A) BT549, (B) HCC1395-OlaR, (C) MDAMB231, and (D) MX1 cells measured by DNA fiber assay mean count of 605 fibres per condition. Dot plots of fork speed (kb/min). Lines indicate median values terquartile range. Kruskal-Wallis test was performed in GraphPad Prism. P<0.0001****, P<0.001***, 1**. (E) Representative images of HCC1395-OlaR DNA fibres at 63x objective. DNA was labelled with magenta) and IdU (green) thymidine analogues. Scale bars represent 10 µm.

### Combined PARPi and LNT1 are synergistic in a xenograft-derived organoid model of TNBC

To evaluate the potential of talazoparib and LNT1 in a more stringent model, we established xenograft-derived organoids (XDOs) (Fig. 5A). Here, we developed organoids from primary tumors in mice that were implanted with the HCC1806 cell line. Once XDOs were established, we characterized the XDOs for their sensitivity to both talazoparib and LNT1 treatment by performing IC_50_ experiments (Fig. 5B, C). We found that, relative to the IC_50_ values we measured across cell lines (Fig. 1A, B), HCC1806 XDOs were more resistant to talazoparib, with an IC50 of 0.11 µM, and slightly resistant to LNT1, with an IC50 of 12.5 µM (Fig. 5C). Thus, based on this high resistance to talazoparib, we considered that HCC1806 XDOs, in contrast to the cell line counterpart, might exhibit synergism between talazoparib and LNT1. Indeed, when treated with the combination, we calculated the CI of talazoparib with LNT1 as 0.80 for HCC1806 XDOs (Fig. 5D). Thus, similar to what we observed in cell lines, we observed that LNT1 synergizes with talazoparib in talazoparib-resistant contexts.

**Figure 5.**
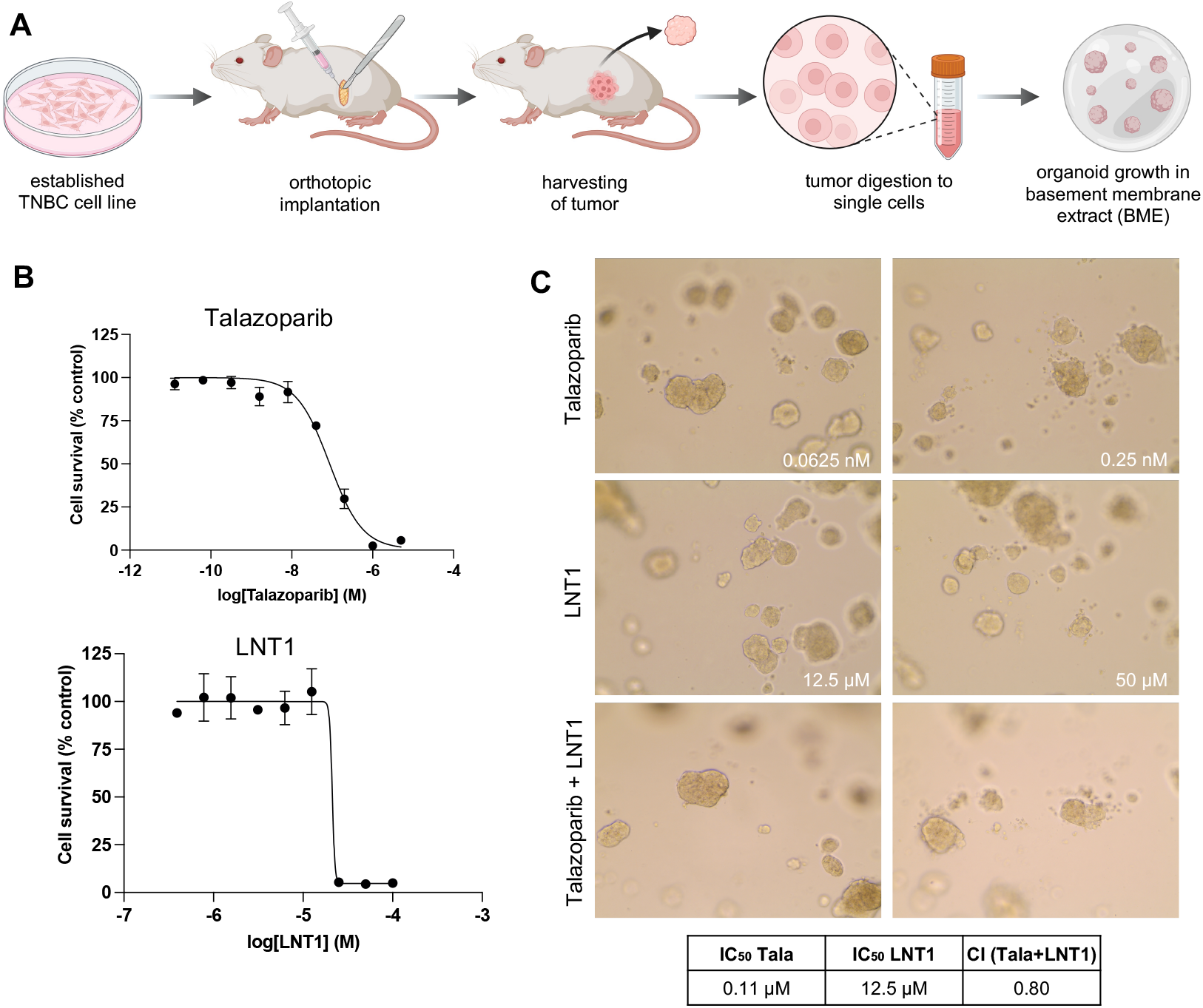
Talazoparib and LNT1 are synergistic in xenograft-derived organoid (XDO) model of TNBC. (A) Shematic representation of generation of XDO. Cells from established TNBC cell lines were implanted e mammary fat pad of mice from which tumors were grown and harvested for organoid development. rs were dissociated to single cells and embedded in basement membrane extract for growth. (B) sentative IC_50_ dose-response curves of talazoparib (top) and LNT1 (bottom) in HCC1806 XDOs. response curves include nine concentrations of talazoparib from 0.013 nM to 5 μM with 1/5 dilutions, concentrations of LNT1 from 0.39 µM to 100 μM with 1/2 dilutions. (C) Representative images of 806 XDO cells treated with talazoparib (top), LNT1 (middle), or the combination (bottom). IC_50_ values azoparib (Tala) and LNT1, and CI (Combination Index) for the combination are indicated below.

### Genetic determinants of response to LNT1 and synergy with talazoparib

To better understand if FEN1 expression influenced response to LNT1, we evaluated FEN1 across our panel of cell lines (Supplementary Fig. S5). We quantified FEN1 RNA expression using our RNAseq gene expression data, and FEN1 protein expression using immunoblotting (Supplementary Fig. 5A,C,D). We did not find any correlation with FEN1 RNA or protein expression with LNT1 response (Supplementary Fig. 5B,E). Interestingly, in the context of BRCA2-deficient tumors that are PARPi-resistant, loss of Poly (ADP-Ribose) glycohydrolase (PARG) was synthetically lethal with FEN1 and EXO1^33^. Therefore, we evaluated PARG RNA expression in our TNBC cell lines (Supplementary Fig. 5F). We did not find decreased expression of PARG in HCC1395-OlaR or HCC1395-TalaR cell lines. The lowest PARG expression was in Hs578T, in which the combination was antagonistic, suggesting that low PARG did not sensitize PARPi-resistant cells to FEN1 inhibition.

We then wanted to better understand which features of the PARPi-resistant cell lines might be modulated by FEN1 inhibition, making them sensitive to PARPi. To do so, we performed a pre-ranked Gene Set Enrichment Analysis (GSEA), comparing RNA expression of untreated PARPi-resistant cell lines that became sensitive with LNT1 (BT549, HCC1395-OlaR, HCC1395-TalaR) versus the expression of a PARPi resistant cell that remained resistant despite LNT1 treatment (Hs578T) (Fig. 6). Upregulated pathways include ribosome (Nominal Enrichment Score (NES) 2.18; False Discovery Rate (FDR) <0.001), Myc targets (NES 1.55; FDR 0.10), mismatch repair (NES 1.53; FDR 0.10), UV response up (NES 1.45; FDR 0.13), and chemokine signalling (NES 1.31; FDR 0.23) (Fig. 6B). Downregulated pathways include epithelial-mesenchymal transition (NES -2.32; FDR <0.001) and apoptosis (NES -1.52; FDR 0.11). Hierarchical clustering revealed some similarities and some differences across the PARPi-resistant cell lines (Fig. 6C). This is suggestive that in addition to mismatch repair and DNA damage response genes, FEN1 inhibition is probably implicated in downregulating ribosomal and chemokine signalling genes, which in turn increases sensitivity to PARPi.

**Figure 6.**
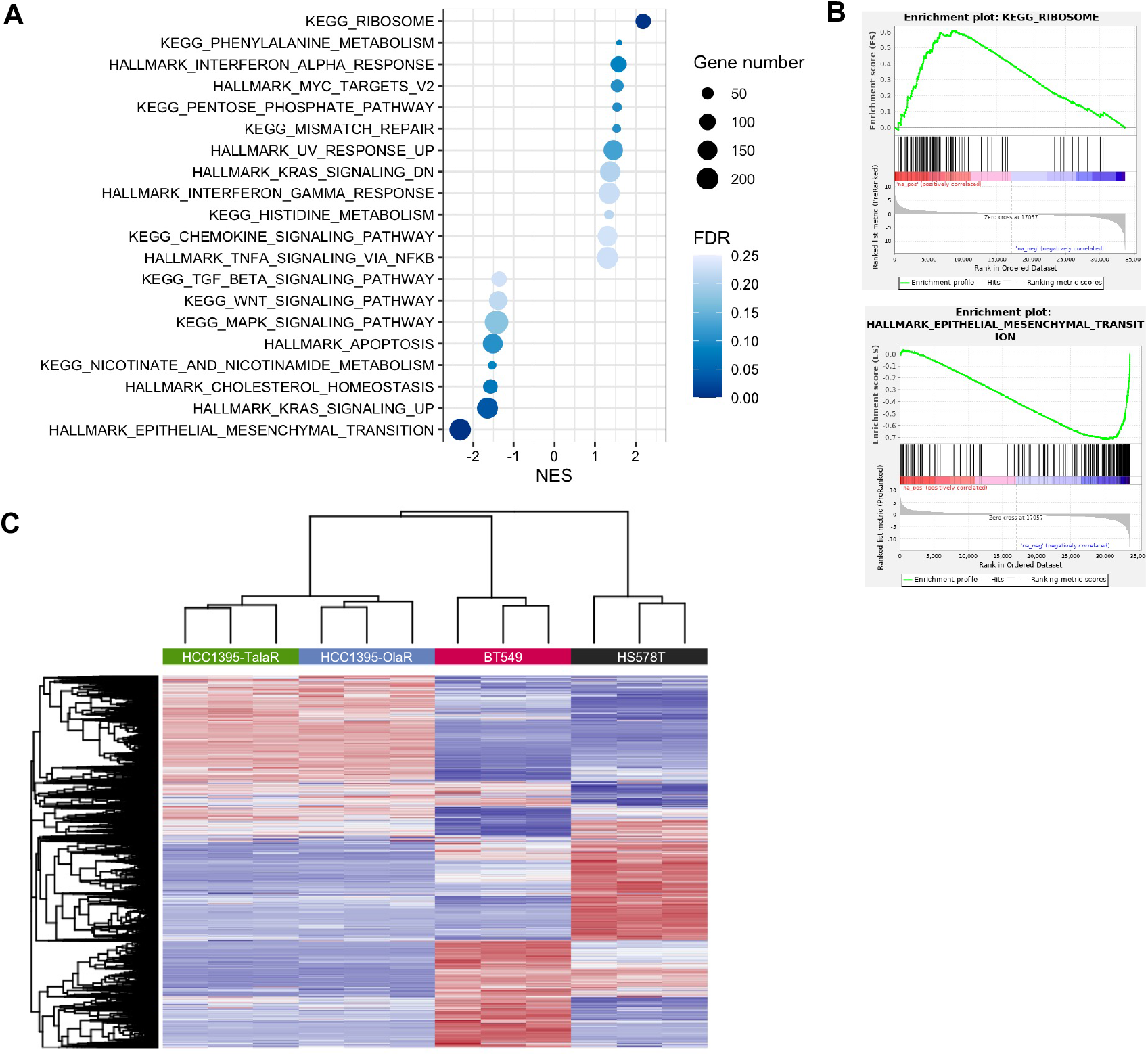
Gene set enrichment analysis (GSEA) of talazoparib-resistant cells reveals PARPi ance mechanisms. (A) Dot plot of the most significantly enriched pathways in GSEA analysis using arks and KEGG datasets, comparing HCC1395-OlaR, HCC1395-TalaR, and BT549 (synergistic) to 8T cells (antagonistic). Pathway names are indicated on the y-axis. Colour intensity indicates the false ery rate (FDR) and dot size indicates the number of genes in each pathway. The x-axis represents al enrichment score (NES), with negative values indicating pathways downregulated in synergistic es and positive values indicating pathways upregulated in synergistic cell lines, relative to HS578T. presentative enrichment plots of the most upregulated (left, ribosome) and most downregulated (right lial-mesenchymal-transition) pathways based on NES, identified by GSEA. (C) Heatmap entation of hierarchical clustering of each cell line with three replicates.

Finally, to investigate the mechanism driving acquired resistance to PARPi in HCC1395 cells, we used a pre-ranked GSEA to compare HCC1395-OlaR and HCC1395-TalaR cells versus HCC1395-Parental. The most upregulated pathway in HCC1395-Ola/Tala-Res was ribosome (NES 2.05; FDR 0.004) (Supplementary Fig. S6). This is suggestive that upregulation of ribosomal genes may be an important mechanism driving intrinsic and acquired resistance to PARPi.

## Discussion

With the increasing utility of PARPi in the clinic^34^, improved strategies are required to tackle resistance to PARPi. We identified several candidate genes from our 63-gene signature that demonstrated great potential if targeted in combination with PARPi. We found that amongst *USP1, FEN1*, and *EXO1*, FEN1 knockdown plus PARPi induced the most apoptosis, thereby providing rationale for selecting FEN1 for further investigation. To our knowledge, we are the first to report the impact of the combination of FEN1 inhibition and PARPi in a panel of patient-derived BRCA^MUT^ and BRCA^WT^ TNBC cell line and organoid models. We showed that the combination of LNT1 and talazoparib demonstrated an additive or synergistic effect in the majority of TNBC cell lines, with synergy demonstrated in PARPi-resistant cell lines, and the strongest synergy demonstrated in a clinically-relevant model, a *BRCA2*-mutant cell line with acquired resistance to olaparib. Furthermore, significant dose reductions for both LNT1 and PARPi suggest the potential for an effective targeted combination with less toxicity.

Our data support three different mechanisms which may be driving the synergistic response in the different cell lines. BT549 is deficient in PTEN which causes significant replication fork stalling^35^, and can explain its lower endogenous fork speed. Together with a mutation in *ATR*^*22*^, BT549 was the only cell line to increase fork speed by targeting FEN1 alone. In this context, unligated Okazaki fragments can promote PARP-1 activity^36^, allowing PARPi to strongly increase fork speed and DNA damage. While we did not observe an impact of the combination on cleaved-PARP expression, this is similar to what we previously observed with the synergistic combination of PARPi and carboplatin in BT549, since there may be other pathways through which PARPi can induce cell death^7,22^.

Second, we observed that the DNA damage response is enhanced when PARPi and LNT1 are combined in all HCC1395 cells. Indeed, FEN1 was shown to be implicated in microhomology-mediated-end-joining (MMEJ) as a collateral repair pathway in the context of homologous recombination deficiency, and thus a synthetically lethal target in *BRCA2*-mutant cells^19^. In response to UV irradiation, phosphorylated FEN1 migrates from the nucleolus, where it is accumulated, to the nuclear plasma to rescue stalled DNA replication forks, and translocated back to nucleolus upon DNA repair^37^. Hence, it is probable that the sustained DNA damage induced by PARPi depletes nucleolar FEN1, thereby amplifying the DNA damage response and cell death.

Finally, there may be another mechanism which can explain the synergy amongst the PARPi-resistant cell lines. We demonstrated that mismatch repair and ribosomal pathways were upregulated in PARPi-resistant cell lines, similar to what was previously reported in mouse mammary cancer cell lines and tumors^33,38^. While PARP-1 binds to small nucleolar RNAs that stimulate PARP-1 activity, PARPi impedes ribosomal DNA transcription and ribosome biogenesis^39^. *FEN1* knockdown was also shown to be involved in downregulating ribosomal RNA processing and ribosome biogenesis in HEK293T cells^40^. Therefore, it is plausible that a 2-pronged attack on ribosome biogenesis may be an important mediator of the synergy observed with talazoparib and LNT1.

We also demonstrated that FEN1 overexpression was associated with a poorer survival in breast cancer patients. These results are similar to what was previously reported in breast cancer and other cancer types^41-44^. FEN1 was shown to be overexpressed in metastatic breast tissue samples in comparison to tumor samples, and in tumor samples in comparison to normal tissue^41^. However, we did not observe a correlation with FEN1 RNA or protein expression and response to LNT1. FEN1 regulation is complex, with multiple interacting proteins, post-translational modifications, and localization in various subcellular compartments^45^. Thus, total RNA or protein expression may not be the best predictive biomarker.

Taken together, targeting FEN1 with PARPi is an effective combination strategy to increase replication fork speed, DNA damage, cell cycle arrest, and cell death. Importantly, this combination approach demonstrated synergy in both intrinsically and acquired PARPi-resistant cells. Since current FEN1 inhibitors have been difficult to use in-vivo due to low potencies and pharmacokinetic profiles^46^, our study supports further investigation in the development of new anti-FEN1 compounds. Targeting FEN1 and PARP-1/2 demonstrates great potential to provide a new targeted combination approach for TNBC patients, leading to decreased toxicity and improved survival.

## Additional Information

## Acknowledgements

We thank the Cellular Imaging, Cellular Physiology, and Animal Platforms of the Centre de Recherche de Centre hospitalier de l’Université de Montréal (CRCHUM). We thank Raphaelle Lambert and Patrick Gendron from the Genomics and Bioinformatics Facility of the Institut de Recherche en Immunologie et Cancérologie (IRIC) for the RNA-Seq analysis. Schematic diagrams were created with BioRender.com.

## Authors’ contributions

Conceptualization, S.H.; Methodology, E.F., M.I.F., S.H.; Investigation, E.F., A.C., D.A., M.I.F.; Visualization, M.I.F, Formal Analysis, E.F., A.C., M.I.F; Writing – Original Draft, M.I.F; Writing – Review & Editing, M.I.F. and S.H.; Funding Acquisition, S.H.; Resources, S.H..; Project administration, S.H., Supervision, S.H. All authors have seen and accepted the final version of the manuscript.

