## Supplementary Tables and Methods for "Targeting FEN1 to enhance efficacy of PARP inhibition in triple-negative breast cancer"

### Supplementary Methods Table S1. Key Resources Table

| Reagent | Source | Identifier |
|---|---|---|
| ***Antibodies*** | | |
| Cleaved-PARP | Cell Signaling Technology | Cat #9546 |
| 53BP1 | Novus Biologicals | Cat #NB100-904 |
| Alexa 647 donkey anti-rabbit | Life Technologies | Cat #A31573 |
| Alexa 488 donkey anti-mouse | Life Technologies | Cat #A21202 |
| HCS Nuclear Mask | Life Technologies | Cat #H10325 |
| gH2AX-AlexaFluor 647 | BioLegend | Cat #613408 |
| Cleaved-caspase 3-Pacific Blue | Cell Signaling Technology | Cat #8788 |
| Rat anti-BrdU (clone BU1/75 (ICR1)) | Abcam | Cat #ab6326 |
| Mouse anti-BrdU [clone B44] | BD Biosciences | Cat #347580 |
| Anti-Rat AlexaFluor 568 | Invitrogen | Cat #A11077 |
| ***siRNAs*** | | |
| ON-TARGETplus Non-targeting Pool, 5 nmol | Horizon | Cat #D-001810-10-05 |
| ON-TARGETplus Human CHEK1 (1111) siRNA - SMARTpool | Horizon | Cat #L-003255-00-0005 |
| ON-TARGETplus Human BRCA1 (672) siRNA - SMARTpool | Horizon | Cat #L-003461-00-0005 |
| ON-TARGETplus Human BARD1 (580) siRNA - SMARTpool | Horizon | Cat #L-003873-00-0005 |
| ON-TARGETplus Human USP1 (7398) siRNA - SMARTpool | Horizon | Cat #L-006061-00-0005 |
| ON-TARGETplus Human BUB1 (699) siRNA - SMARTpool | Horizon | Cat #L-004102-00-0005 |
| ON-TARGETplus Human FEN1 (2237) siRNA – SMARTpool | Horizon | Cat #L-010344-00-0005 |
| ON-TARGETplus Human EXO1 (9156) siRNA - SMARTpool | Horizon | Cat #L-013120-00-0005 |
| ON-TARGETplus Human RRM2 (6241) siRNA - SMARTpool | Horizon | Cat #L-010379-00-0005 |
| ***Chemicals, peptides, and recombinant proteins*** | | |
| Talazoparib | Selleckchem | Cat #S7048 |
| Olaparib | Selleckchem | Cat #S1060 |
| LNT1 | Tocris Bioscience | Cat #6510/5 |
| Propidium Iodide | Invitrogen | Cat #P3566 |
| Chloro-2'-deoxyuridine (CldU) | Sigma-Aldrich | Cat #C6891 |
| 5-Iodo-2'-deoxyuridine (IdU) | Sigma-Aldrich | Cat #I7125 |
| R-spondin 1 | PeproTech | Cat #120-38-100UG |
| Noggin | PeproTech | Cat #120-10C-100UG |
| Nicotinamide | ThermoFisher | Cat #A15970.22 |
| SB202190 | StemCell Technologies | Cat #202190 |
| ***Critical Commercial Assays*** | | |
| Cell lysis buffer | Cell Signalling Technology | Cat #9803 |
| PhosSTOP | Roche | Cat #4906845001 |
| Cultrex Reduced Growth Factor Basement Membrane Extract | R&D Systems | Cat. #3433-005-01 |
| ProLong™ Gold antifade Mountant | Invitrogen | Cat #P36930 |
| Pierce BCA Protein Assay Kit | ThermoFisher | Cat #23225 |
| Mycoalert mycoplasma detection kit | Lonza | Cat #LT07 |
| Lipofectamine RNAiMAX transfection reagent | ThermoFisher | Cat #13778150 |
| Opti-MEM | ThermoFisher | Cat #31985062 |
| CellTiter-Glo 3D Cell Viability Assay | Promega | Cat #G9682 |
| ***Experimental models: Cell lines*** | | |
| MDAMB231 | JW Gray Lab | N/A |
| Hs578T | JW Gray Lab | N/A |
| SUM149PT | JW Gray Lab | N/A |
| HCC1395 | JW Gray Lab | N/A |
| HCC1806 | JW Gray Lab | N/A |
| MX1 | JW Gray Lab | N/A |
| MDAMB436 | ATCC | Cat #HTB-130 |
| BT549 | ATCC | Cat #HTB-122 |
| ***Experimental models: Mice*** | | |
| NOD.Cg-Prkdcscid Il2rgtm1Wjl/SzJ | The Jackson Laboratory | Cat# 005557 |
| ***Software and algorithms*** | | |
| GraphPad PRISM 10 | GraphPad software | https://www.graphpad.co |

| | | |
|---|---|---|
| | | m/scientific-software/prism/ |
| STATA SE, Version 15 | StataCorp LLC | https://www.stata.com/ |
| FlowJo | BD Biosciences | https://www.flowjo.com/ |
| Compusyn | ComboSyn Incorporated | https://www.combosyn.com |
| ImageJ | NIH | https://imagej.nih.gov/ij/ |
| Harmony High Content Imaging and Analysis Software | Perkin Elmer | https://www.perkinelmer.com/product/harmony-4-8-office-hh17000001 |
| GenePattern, Version 3.9.11 | National Cancer Institute's Informatics Technology for Cancer Research program and the National Institute of General Medical Sciences | https://cloud.genepattern.org/gp/pages/index.jsf |

**Supplementary Table S2.** List of antibodies used for western blotting.

| Antibody | Company | Cat. Number | Dilution | Blocking buffer |
|---|---|---|---|---|
| Anti-BARD1 | Santa Cruz Biotechnology | Sc-74559 | 1:500 | 5% Skim milk - TBST |
| Anti-β-actin | ThermoFisher | AM4302 | 1:5000 | 5% Skim milk - TBST |
| Anti-BRCA1 | Novus Biologicals | NB-100-404 | 1:500 | 5% Skim milk - TBST |
| Anti-BUB1 | ThermoFisher | A300373A-M | 1:1000 | 5% Skim milk - TBST |
| Anti-EXO1 | Abcam | ab95068 | 1:2000 | 5% Skim milk - TBST |
| Anti-FEN1 | Abcam | ab109132 | 1:1000 | 5% Skim milk - TBST |
| Anti-RRM2 | Abcam | ab172476 | 1:200 | 5% Skim milk - TBST |
| Anti-USP1 | Cell Signaling Technology | 8033 | 1:1000 | 5% Skim milk - TBST |
| Anti-mouse HRP | Biorad | 1706516 | 1:2000 | 5% Skim milk - TBST |
| Anti-rabbit HRP | Biorad | 1706515 | 1:2000 | 5% Skim milk - TBST |

**Supplementary Table S3.** Drug concentrations used in Figure 3A-B.

| Cell line | [Talazoparib] (μM) | [LNT1] (μM) |
|---|---|---|
| BT549 | 0.020 – 0.63 | 1.6 - 50 |
| HCC1395-Parental | $3.1 \times 10^{-5}$ – 0.0010 | 1.6 - 50 |
| HCC1395-OlaR | 0.0094 – 0.30 | 3.1 - 100 |
| HCC1395-TalaR | 0.0016 – 0.050 | 3.1 - 100 |
| HCC1806 | $3.1 \times 10^{-5}$ – 0.0010 | 0.5 - 16 |
| MDAMB231 | 0.0050 – 0.16 | 1.9 - 60 |

### Supplementary Methods

*siRNA transfections*

For each human gene target, we used ON-TARGET-plus SMARTpool siRNA (4 siRNA targets/gene) with a non-target pool as control (Dharmacon). MDAMB231 cells were transfected with siRNA using Lipofectamine RNAiMAX transfection reagent (Cat. #13778150) in 6-well plates. 60 pmol of siRNA was diluted in 500 µL of Opti-MEM (Cat. #31985062), then 5 µL of lipofectamine was added. The combination was vortexed then incubated at room temperature for 10 to 20 minutes. Finally, 500 µL of the siRNA/lipofectamine suspension was added to cells with 2.5 mL of culture media. After 24 hours, the media was changed. Cells were then treated with 0.25 µM talazoparib or DMSO control for 72 hours, then harvested for flow cytometry.

*Chemosensitivity assays*

10-day chemosensitivity assays were performed as we previously described[9,26]. Cells were treated with 9 concentrations of talazoparib (5-fold dilutions) or LNT1 (2-fold dilutions). The maximum concentrations were 5 µM and 100 µM for talazoparib and LNT1, respectively. Dose-response curves were created in GraphPad Prism 10 software and used to derive $IC_{50}$ values. IC50 values were obtained from 2-5 replicate assays.

*High-content imaging*

Fixed and stained cells were imaged on the Operetta CLS (PerkinElmer) with 20X objective and filters for Alexa 488, Alexa 647, and DAPI. 37 images were scanned per well and image analysis was performed with Harmony High-Content Imaging and Analysis Software (version 5.2, PerkinElmer).

*Combination index values and Immunofluorescence staining*

Chemosensitivity treatments were performed as we previously described[26]. Cells were treated with six concentrations of talazoparib, LNT1, or the combination, with 1/2 drug dilutions, and concentrations centred around the $IC_{50}$ for each drug for each specific cell line, using the 10-day chemosensitivity assay. Cells were fixed and stained with DAPI alone and/or 53BP1 or cleaved-PARP for detection via immunofluorescence. Combination Index (CI) and Dose Reduction Index (DRI) values were calculated by the Chou-Talalay method using Compusyn software (ComboSyn Incorporated, Paramus, NJ, USA) at Fa = 0.50. CI values between 0.9 and 1.10 are considered nearly additive; 0.85–0.7 demonstrate moderate synergism, and 0.3–0.7 indicate synergism[27].

*DNA fibre assay and flow cytometry*

Cells in 6-well plates were treated with talazoparib, LNT1, or the combination for 72 hours, at different combinations depending on sensitivity to each drug. DNA fibre assays and flow cytometry experiments were then performed exactly as described[26].
