## Supplementary Figures for "Targeting FEN1 to enhance efficacy of PARP inhibition in triple-negative breast cancer"

#### Supplementary Figure S1

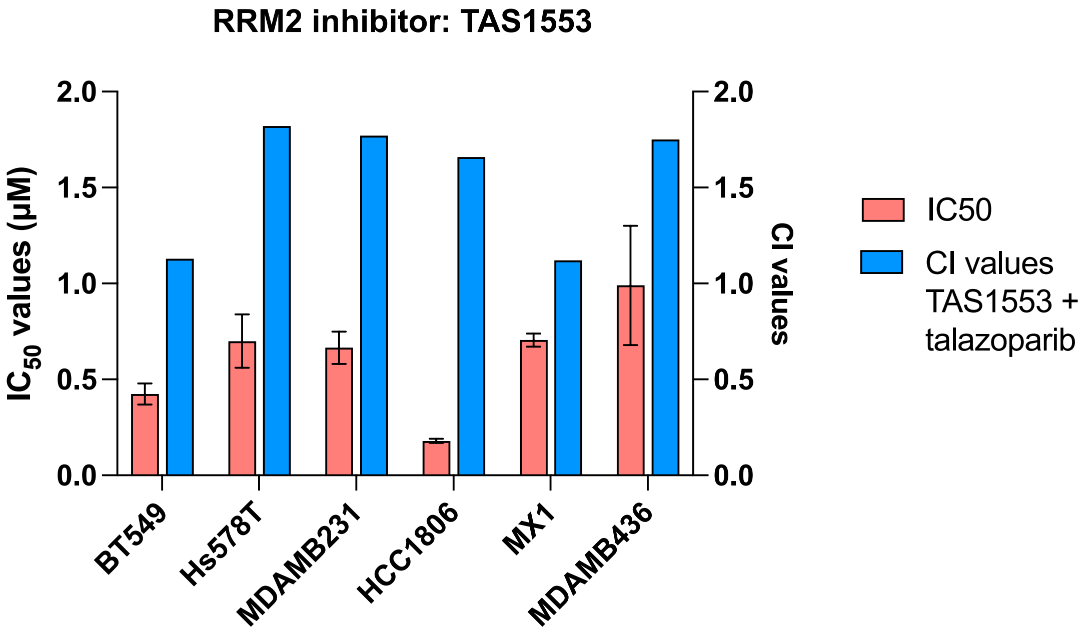


**Figure S1. RRM2 inhibition does not sensitize cells to talazoparib.** $IC_{50}$ (pink) and combination index (CI, blue) values for the RRM2 inhibitor TAS1553 in six TNBC cell lines.

### Supplementary Figure S2

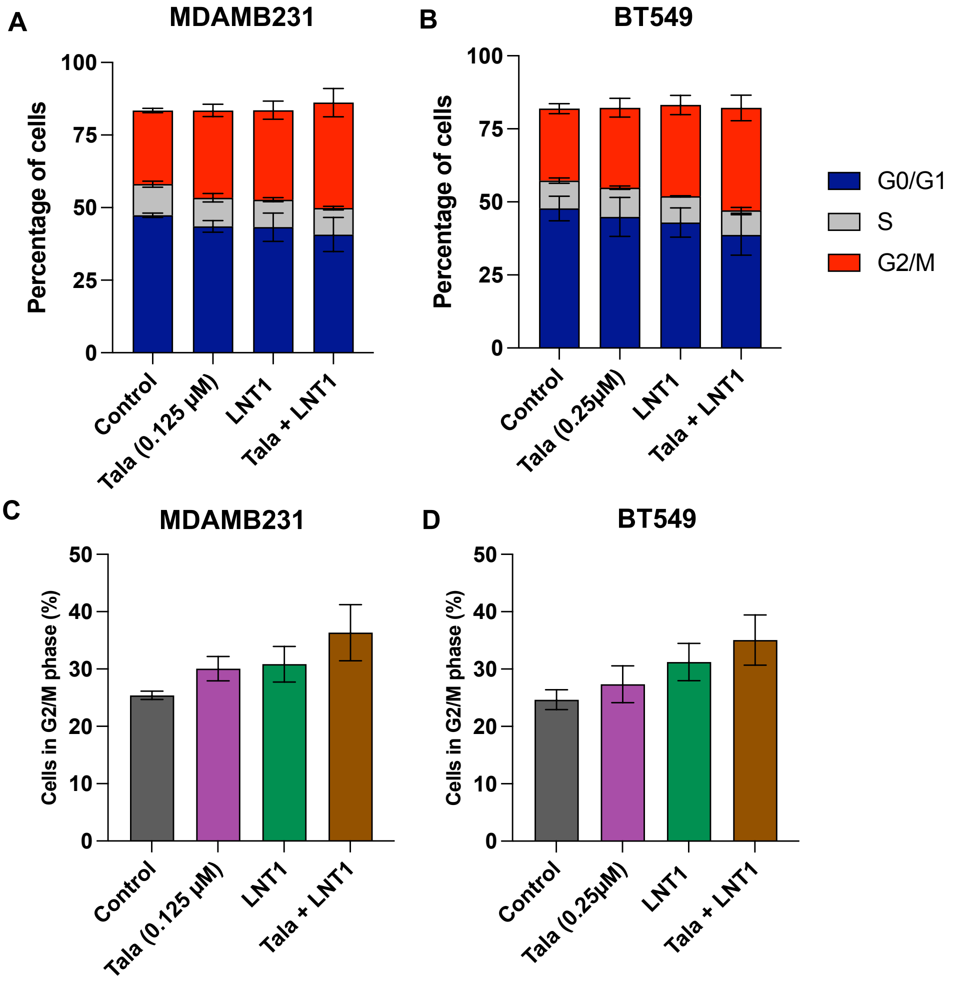


**Figure S2. Impact of 72-hour treatments on cell cycle.** Impact of talazoparib or LNT1 treatment on cell cycle changes are shown in (A,C) MDAMB231 and (B,D) BT549 cells.

### Supplementary Figure S3

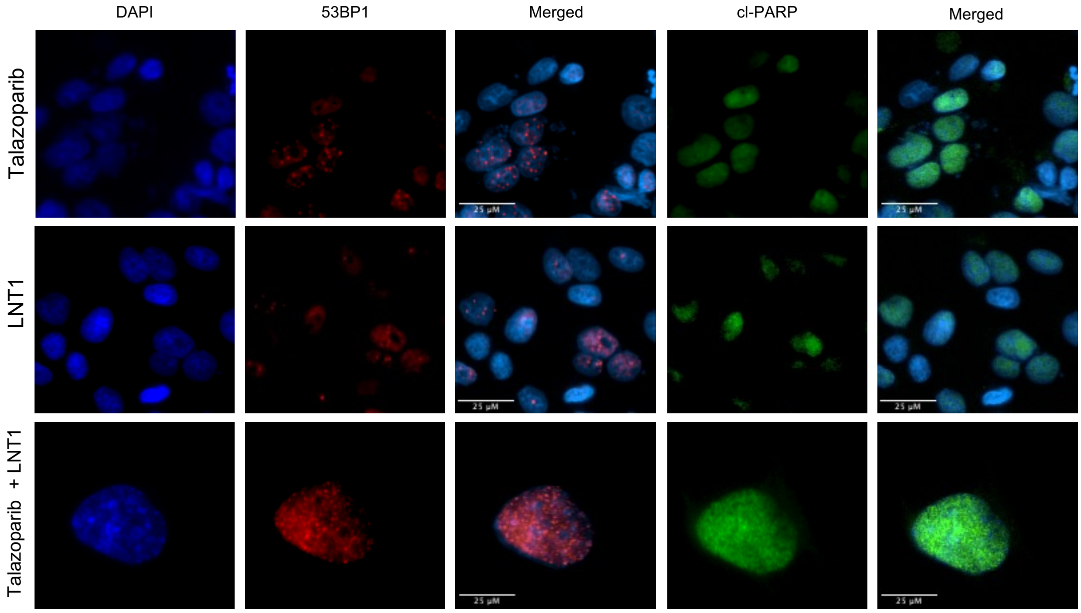


**Figure S3. 53BP1 foci and cleaved (cl)-PARP expression in HCC1395-OlaR cells.** Representative single channel images of immunofluorescence. Far left column shows nuclei stained with HCS nuclear mask, middle left column indicates 53BP1 foci, and middle column are merged 53BP1/HCS nuclear mask. Middle right column indicates cl-PARP expression, and far right column are the merged cl-PARP/HCS nuclear mask images. Scale bars indicate 25 μm.

### Supplementary Figure S4

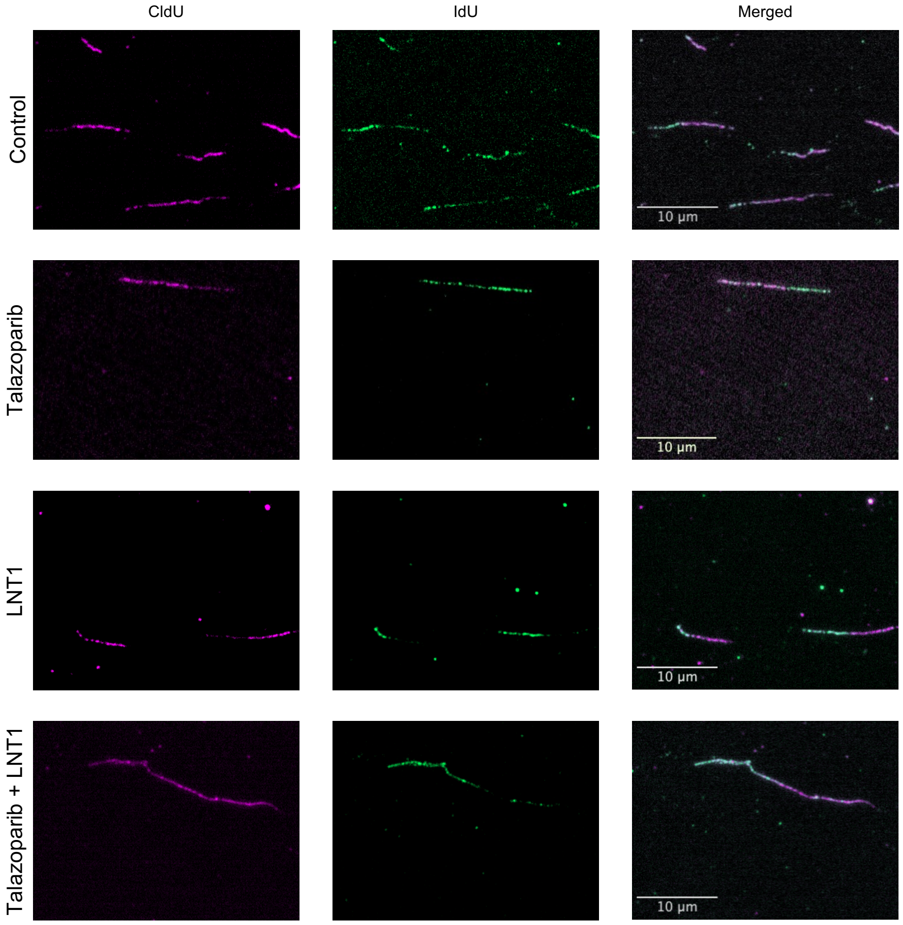


**Figure S4. DNA fiber images of HCC1395-OlaR cells treated with talazoparib or LNT1 as single-agents or in combination.** Representative single-channel images of CldU in magenta (left), IdU in green (middle), and merged images (right). Scale bars refer to 10 microns.

### Supplementary Figure S5

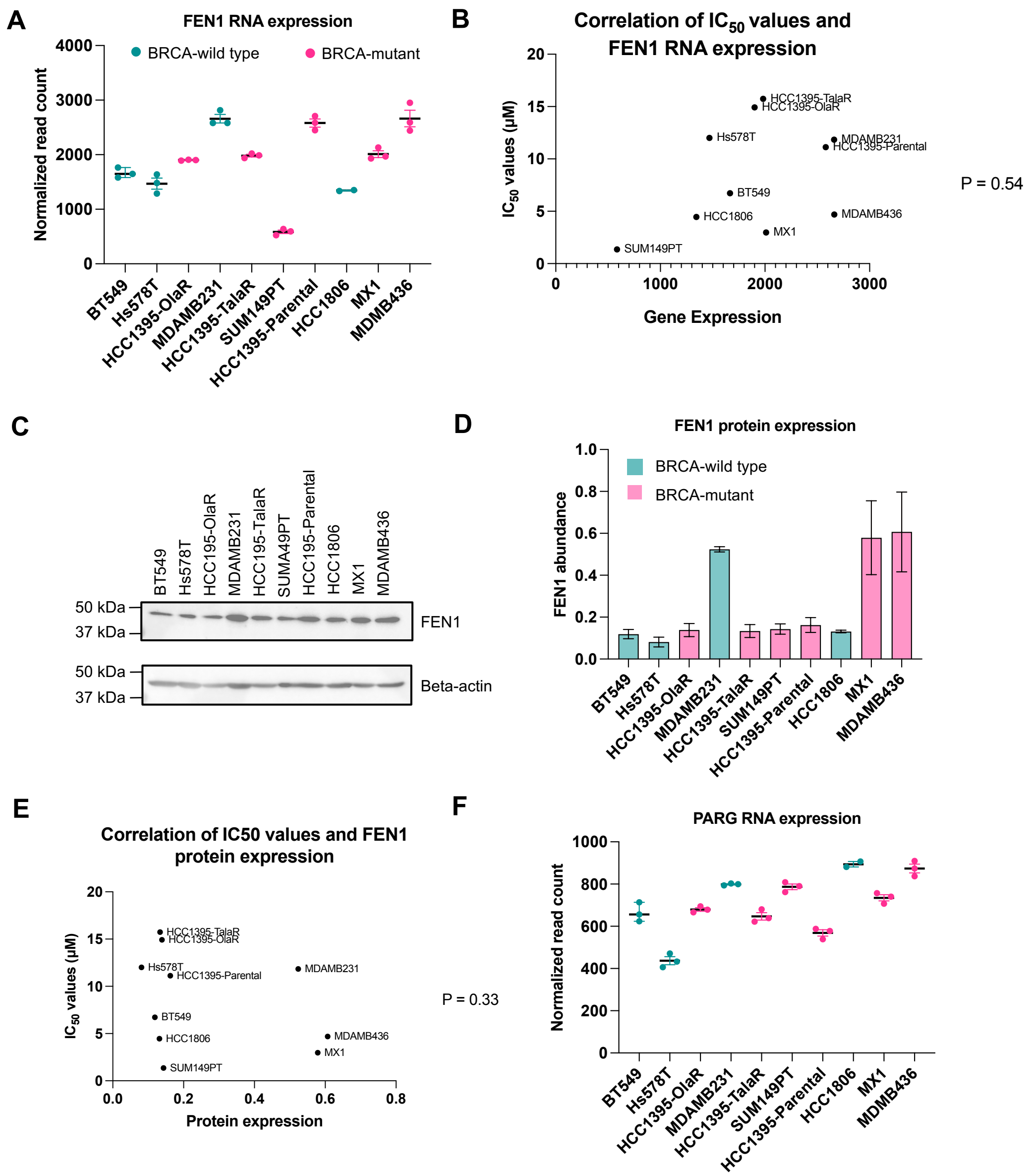


**Figure S5. FEN1 expression in TNBC.** (A) FEN1 expression levels across TNBC cell lines, measured by RNA-seq. (B) Correlation of FEN1 abundance measured by RNA-seq with LNT1 $IC_{50}$ values. (C) Western blot and (D) quantification of FEN1 in TNBC cell lines. (E) Correlation of FEN1 abundance measured by western blot with LNT1 $IC_{50}$ values. (F) PARG expression levels across TNBC cell lines, measured by RNA-seq. Teal green bars/dots represent $BRCA^{WT}$ cell lines, and pink bars/dots represent $BRCA^{MUT}$ cell lines.

### Supplementary Figure S6

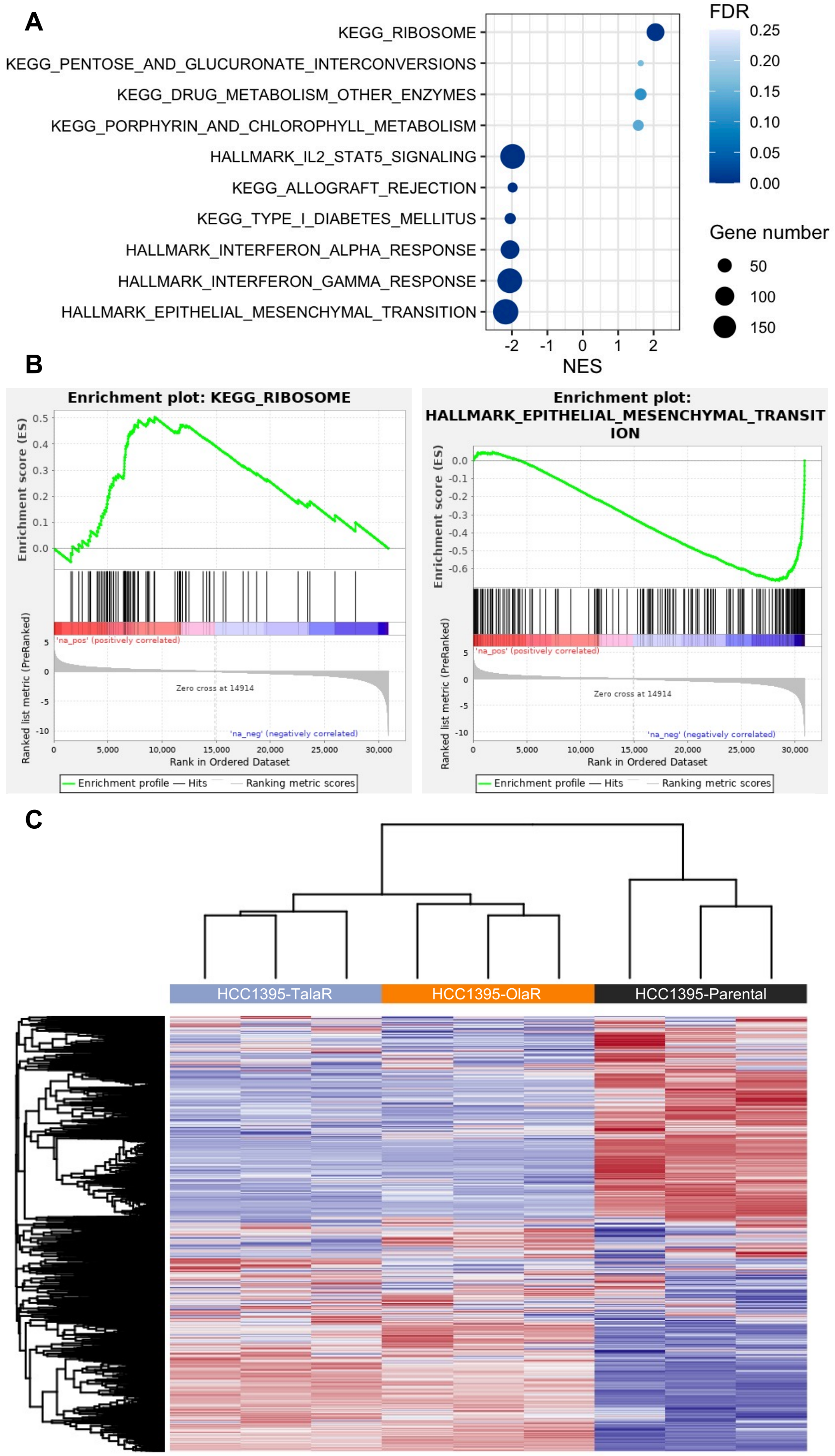

**Figure S6. Gene set enrichment analysis (GSEA) of HCC1395 cells reveals pathways associated with PARPi resistance.** (A) Dot plot of the most significantly enriched pathways in GSEA analysis using Hallmarks and KEGG datasets, comparing HCC1395-OlaR and HCC1395-TalaR to HCC1395-Parental cells. Pathway names are indicated on the y-axis. Colour intensity indicates the false discovery rate (FDR) and dot size indicates the number of genes in each pathway. The x-axis represents nominal enrichment score (NES), with negative values indicating pathways downregulated in HCC1395-OlaR and HCC1395-TalaR and positive values indicating pathways upregulated in HCC1395-OlaR and HCC1395-TalaR, relative to HCC1395-Parental. (B) Representative enrichment plots of the most upregulated (left, ribosome) and most downregulated (right, epithelial-mesenchymal-transition) pathways based on NES, identified by GSEA (C) Heatmap of hierarchical clustering of triplicate cell lines: HCC1395-Parental (black), HCC1395-OlaR (orange), and HCC1395-TalaR (blue).
